# Identification and organization of a postural anti-gravity module in the cerebellar vermis

**DOI:** 10.1101/2023.10.10.561555

**Authors:** Aurélien Gouhier, Vincent Villette, Benjamin Mathieu, Annick Ayon, Jonathan Bradley, Stéphane Dieudonné

## Abstract

The cerebellum is known to control the proper balance of isometric muscular contractions that maintain body posture. Current optogenetic manipulations of the cerebellar cortex output, however, have focused on ballistic body movements, examining movement initiation or perturbations. Here, by optogenetic stimulations of cerebellar Purkinje cells, the output of the cerebellar cortex, we evaluate body posture maintenance. By sequential analysis of body movement, we dissect the effect of optogenetic stimulation into a directly induced movement that is then followed by a compensatory reflex to regain body posture. We identify a module in the medial part of the anterior vermis which, through multiple muscle tone regulation, is involved in postural anti-gravity maintenance of the body. Moreover, we report an antero-posterior and medio-lateral functional segregation over the vermal lobules IV/V/VI. Taken together our results open new avenues for better understanding of the modular functional organization of the cerebellar cortex and its role in postural anti-gravity maintenance.

**Highlights:**

- Vermal Purkinje cell activation elicits a graded postural collapse in the standing mouse
- The collapse triggers a secondary composite postural reflex
- An identified cerebellar module is involved in postural anti-gravity tone maintenance
- The anti-gravity function is somatotopically organized within this module

## INTRODUCTION

Posture is defined by the arrangement of body parts in space resulting from muscle contractions (Horak 2006). Mechanisms of postural control obviously operate during movements, but as well by giving anti-gravity support to the body during maintenance of static posture (Massion and Dufosse 1988; Horak and Macpherson 1996; Horak 2006). Calculation of the motor commands required to achieve the appropriate tone of effector muscles for static posture is a subconscious process in which the anterior cerebellar vermis plays a crucial role, as lesions of this region cause atonia of the axial muscles as well as stance and gait deficits (Chambers and Sprague 1955a; 1955b; Joyal et al. 1996). While optogenetic tools have been used to directly probe the functional output of some cerebellar regions, a systematic mapping has yet to be produced. Depending on the location and extent of the cortical region targeted*, in vivo* optogenetic manipulation of the cortical cerebellar Purkinje cells (PC) output can induce either discrete movements (Heiney et al. 2014), whole limb motions (Lee et al. 2015), locomotor and rhythmic movement perturbations (Hoogland et al. 2015; Sarnaik and Raman 2018; Gao et al. 2018; Gaffield, Sauerbrei, and Christie 2022), or even whole-body twitches (Witter et al. 2013). Vermal stimulations, however, have thus far failed to reveal a tonic role of the cerebellum in anti-gravity postural control (Witter et al. 2013; Hoogland et al. 2015).

The cerebellum is well-known to be organized in cortico-nuclear-olivo loop modules, each encompassing a subregion of the inferior olive (IO), which projects to a translobular parasagittal band of PCs via the climbing fibers (CFs), and to a subregion of the deep cerebellar nuclei (DCN) supposed to be interconnected in closed loops (Groenewegen and Voogd 1977; Groenewegen, Voogd, and Freedman 1979; Sugihara and Shinoda 2007; Sugihara 2011; Ruigrok 2011). How the DCN outputs relate functionally to this modular organization has been the subject of attention (Fujita, Kodama, and Du Lac 2020; Heiney, Wojaczynski, and Medina 2021; Wang et al. 2023). Recently, specific regions of the brain that are related to several generic posture-motor functions were shown to receive projections from genetically identified subdivisions of the medial nucleus, which are also innervated by PCs of the vermis and the hemispheres (Fujita, Kodama, and Du Lac 2020). The anterior vermis could be subdivided in several broad modules, which may correspond to the ones previously identified by axonal morphology (Voogd and Glickstein 1998), and zebrin immunolabeling (Hawkes, Colonnier, and Leclerc 1985; Sugihara 2011). Several lines of evidence, however, have pointed to a finer modular organization of the vermal PC output. Cutaneous and nerve stimulations have suggested that cerebellar modules could be further divided into microzones of 100-200 µm based on their CF receptive fields (Andersson and Oscarsson 1978; Jörntell et al. 2000). Functional microzones have also been identified by large-scale imaging and correlation of CF discharge in the dorsal cerebellar vermis (Ozden et al. 2009; Mukamel, Nimmerjahn, and Schnitzer 2009; Kostadinov, Beau, Blanco-Pozo, et al. 2019). Furthermore, based on retrograde tracings, it was proposed that thin bands of PCs could act to control single muscles (Ruigrok et al. 2008; Ruigrok 2011). Thus, the functional organization of the cortical cerebellar output remains ineffectually characterized.

In this study, using spatially confined optogenetic stimulations in freely-moving mice expressing the actuator ChR2 specifically in PCs (Chaumont et al. 2013), we aimed at investigating the functional micro-organization of the vermal lobules IV/V/VI output. We found that the vermis is indeed involved in postural maintenance, and more specifically in anti-gravity support. Optogenetic stimulations elicited an initial perturbation of postural maintenance in the quiet animal. This was followed by a sequential postural reflex involving several parts and muscles of the animal’s body. Post-stimulation, we then observe a rebound contraction in the affected muscles. Using a custom fiber array to map the output of the vermis, we reveal that the identified postural function is encoded at the scale of the previously identified A zone, encompassing a large part of the vermal region. Moreover, our data strongly suggest an antero-posterior and medio-lateral functional segregation across the lobules IV/V/VI. Together, our results indicate that computations performed by PCs in the anterior vermis likely involve multi-segmental control of the body to ensure postural anti-gravity maintenance.

## EXPERIMENTAL PROCEDURES

### Mice

We used male and female mice heterozygous for L7-ChR2-eYFP of 6-16 weeks of age (Chaumont et al. 2013). Mice were housed in standard conditions (12-hour light/dark cycles, light on at 7 a.m., with water and food ad libitum). All protocols adhered to the guidelines of the French National Ethic Committee for Sciences and Health report on Ethical Principles for Animal Experimentation in agreement with the European Community Directive 86/609/EEC under agreement #12007.

### Implantation procedure

Mice were anesthetized with intraperitoneal injection of ketamine (100 mg/kg) and xylazine (20 mg/kg, Centravet) and received preoperative analgesic (buprenorphine, 0.1 mg/kg). Mice were then fixed on a stereotaxic frame and a ∼6*4 mm craniotomy was performed over the vermal region of the cerebellar lobule IV/V/VI. The bottom part (head plate) of a custom designed 3-D printed implant was aligned with the lambda-bregma axis using a custom-made head plate holder and fixed on the skull with a layer of dental cement (Superbond). Mice were then temporarily detached from the stereotaxic frame and reattached using custom head plate holding bars. A custom designed laser-cut coverslip was then lowered on top of the targeted vermal area while vacuum-sucked onto a rectified hollow rod mounted on the stereotaxic apparatus. This protocol ensured that the coverslip laid flat on the brain and was parallel to the head plate surface, so that in subsequent in vivo experiments the optic fiber would lie flat on the coverslip. A 300-600 µm excess z drive was performed, thus applying pressure on the brain with the coverslip, to ensure that there would be no space between the coverslip and the cerebellum, which could enhance bone regrowth and brain movement. The coverslip was then secured with dental cement (Superbond). Finally, the upper part of the implant was screwed on the bottom part using two M1 screws. Mice were allowed to recover for at least 5 days before handling sessions and housed 1-3 per cage. Meloxicam (10 mg/kg, Centravet) was given daily for 48h after the surgery.

### EMG surgeries

For EMG recordings, mice were additionally implanted with chronic electromyographic electrodes (EMGs). For the animals in which we performed EMGs implantation, the surgery was performed on the same day following the above-described implantation protocol. Mice were supplemented in anesthetics with isoflurane (2%) and prepared for aseptic surgery. An incision was made over the muscle intended for implantation. The skin was separated from underlying fascia. For splenius muscle implantation, the overlying trapezius muscle was gently dilacerated in the antero-posterior orientation with soft tip tweezers to allow for muscle access. Each pair of electrodes was funneled through subcutaneously from the incision to the back of the head plate implant, where the connector pins were cemented. Each wire was then passed through a small surgical needle (Kalt suture needles size 3, Fine Science Tools #12050-03). The needle was passed through the intended muscles perpendicular to the muscle fibers until the proximal knot pressed against one side of the muscle. Two electrodes were placed in each muscle at a distance of approximately 1 mm. Individual knots were made with the distal ends of the electrodes against the other side of the muscle. The excess of wire was finally cut approximately 2 mm from the muscle surface. After ensuring that the electrodes were in place, incisions were closed with nylon sutures. Mice received postoperative analgesic (buprenorphine, 0.1 mg/kg) and meloxicam (10 mg/kg, Centravet) was given daily for 48h after the surgery.

### Construction of EMG electrodes

We built EMG electrodes based on a previously described procedure (Tysseling et al. 2013). A section of electrode wire (A/M systems, multi stranded stainless steel, Teflon insulated, #793200) was cut whose length depended on the distance between the head plate implant and the targeted muscle (3.5-7.5 cm range). At the proximal end of the wire, a small amount of wire was stripped and soldered to a 1 mm gold pin (19003-00, Fine Science Tools). At the distal end, five overlapping knots were made on each wire, leaving 2-3 cm of wire excess. Then, approximately 0.5 mm of insulated material was stripped on the distal side of the knot. Electrodes were stored in pairs until muscle implantation.

### Implant design

A custom-made 3D-printed implant was designed using the Solidworks software in order to map the entire accessible vermal area in a single animal. The implant consists of two main parts (Figure 4A). The bottom part is a head plate which is cemented on the skull of the animal. It features two empty volumes in which small sliding parts are introduced. The upper part of the implant lands on the bottom part and can slide on it over 3 mm, allowing the coverage of the entire medio-lateral extent of the vermis. It also features a hole at its center into which the optic fiber passes. When in position, the top part of the implant is tightened to the bottom part by two screws (DIN 84 M1, Micro-Modèle) passing through the small slits. The relative positions of the two parts are measured using landmarks printed at their surface. The 3D impressions were done with a Form 2 SLA 3D printer (Formlabs). Because the antero-posterior position of the implant relative to the brain of the animal was variable, upper implant parts with optic fiber holes at different antero-posterior positions were printed. This allowed not only to properly align the fiber in a reproductible position, but also to perform an antero-posterior mapping, since this part of the implant is removable.

### Optic fiber design

For Figures 1 to 3, regular optic fibers with a 200 µm core diameter (NA = 0.22, MFP _200/220/900-0.22_1m_FCM-MF1.25(F), Doric Lenses) were used. For mapping experiments, a custom patch cord was designed. The distal end of the patch cord, away of the animal, consists of 14 50 µm core optic fibers (NA = 0.22) arranged in a disk (Figure 4A). This end of the patch cord is coupled to a 400 µm optic fiber (NA = 0.22) which serves as a relay for laser illumination. At the proximal end of the patch cord, the optic fibers are arranged in a linear array that covers a surface of 50*960 µm, which allows the stimulation specific PC bands.

### Experimental setup

A custom-designed setup was developed to assess postural behavior during optogenetic stimulations in freely moving mice (Figure 1A). The apparatus consists of a clear glass corridor, 45 cm long, 4.5 cm wide and 20 cm high. The corridor is closed on both sides. Mice were filmed while freely behaving with two high-resolution, highspeed cameras (Basler acA1300-3200um). A mirror (45 cm × 15 cm) was placed below the corridor at an angle of ∼45° to allow simultaneous collection of side and bottom views in order to allow for 3-D analysis of the data (Machado et al. 2015). Lighting consisted of six matrices of 860 nm infrared LEDs (SFH 4557, Osram) which were carefully positioned to maximize contrast and reduce reflection.

**Figure 1.**
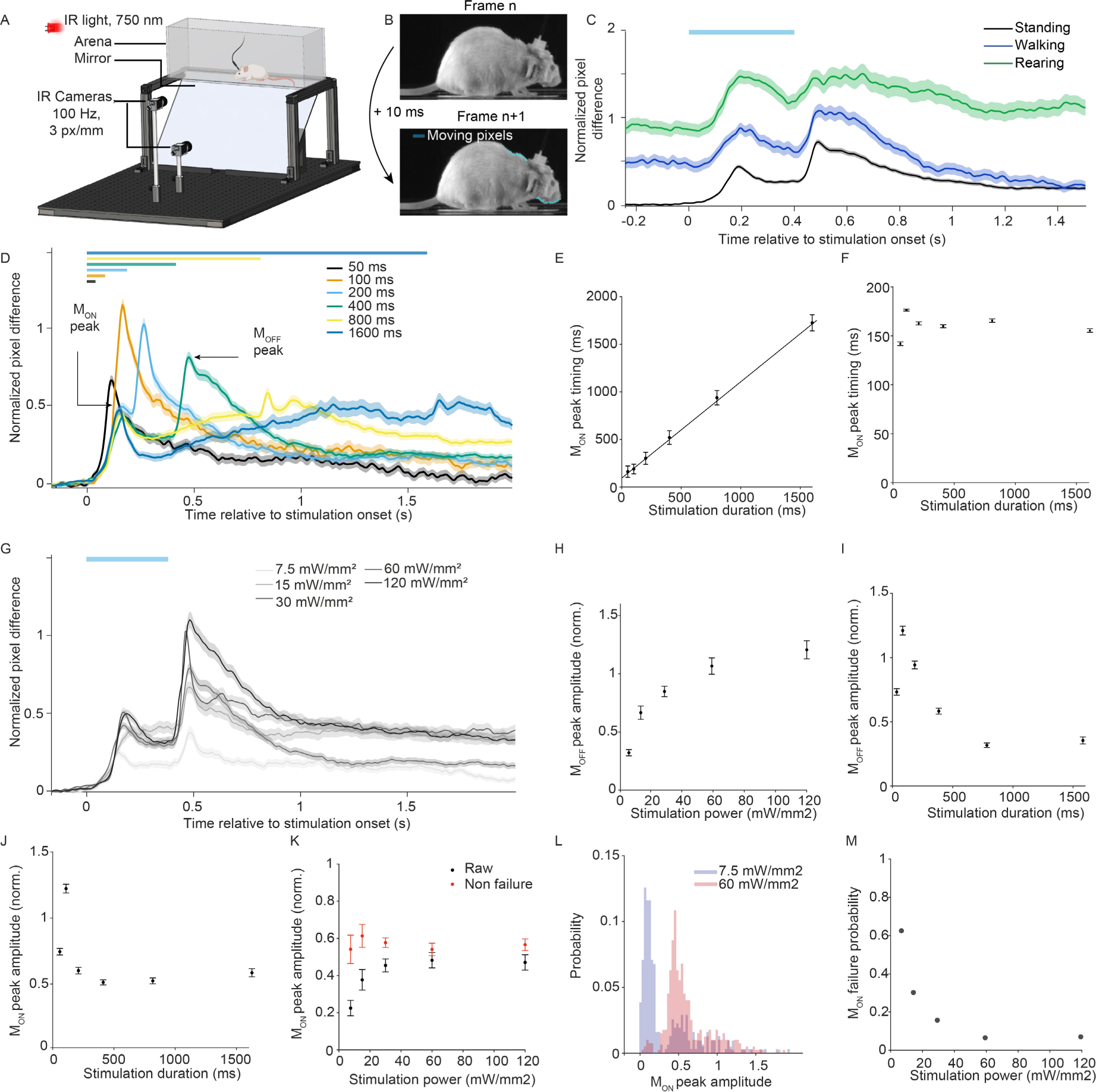
Optogenetic stimulations of the anterior vermal Purkinje cells elicit a movement sequence. **A**. The implanted mice were introduced in a transparent rectangular glass arena with a mirror below at 45° angle. Two high-speed cameras captured side and bottom views at 100 Hz. **B**. Two consecutive frames of the animal side view. Movement was defined as the number of pixels whose values significantly changed between the two frames (light blue areas). **C**. Normalized pixel difference around the optogenetic stimulation for three different behavioral states (see Experimental Procedures). Curves represent mean *±* SEM (n = 8 mice, 30 mW/mm² 400 ms stimulations). **D**. Evolution of the mouse movement with the stimulation duration (n = 8 mice, 50 trials for each mouse and each condition). Curves represent mean *±* SEM. M_OFF_ peak timing plotted against stimulation duration (30 mW/mm² stimulations, R = 0.999, y = 1.02x + 126 ms). Bars represent mean *±* SEM. **F**. M_ON_ peak timing plotted against stimulation duration (30 mW/mm² stimulations, n = 8 mice). **G**. Evolution of the mouse movement with the stimulation power (n = 8 mice, 400 ms stimulations). Curves represent mean *±* SEM. **H.** M_OFF_ peak amplitude plotted against the stimulation power (400 ms stimulations). **I.** M_OFF_ peak amplitude plotted against stimulation duration (30 mW/mm² stimulations). **J.** M_ON_ peak amplitude plotted against stimulation duration (30 mW/mm² stimulations). **K.** Raw and non-failure (see Experimental Procedures) M_ON_ peak amplitude against stimulation power (computed for stimulations longer than 200 ms). **L.** Histograms of M_ON_ peak amplitudes for two stimulation powers (stimulation duration = 400 ms). **M.** M_ON_ failure probability plotted against stimulation power (computed for stimulations longer than 200 ms).

A custom optical setup was placed on top of the arena. It consists of a standard optical breadboard on which a laser (473 nm, LRS-0473, Laserglow Technologies) was aligned with the optic fiber core using a custom-made optical block (Doric Lenses). An acousto-optic tunable filter (AOTFnC-400.650-TN, AAOpto-electronic) was placed on the optical path to allow to trigger optogenetic stimulations. A 5 cm diameter hole was drilled in the breadboard below the optical block which was placed just above the arena. During the experiments, the optic fiber passed directly through the hole and was therefore vertical. This minimized the mechanical constraints on the fiber as well as on the mouse body. To allow for optic fiber rotations, a frictionless rotary joint (Doric Lenses) was used.

### Data collection and acquisition

Mice were handled by the experimenter and allowed to acclimate the setup environment without being introduced into the arena on multiple occasions prior to data collection. Before each experiment, the mouse was head-fixed on top of a non-motorized running wheel. This allowed to unscrew the upper part of the implant and carefully clean the glass coverslip. The upper part of the implant was then positioned to the desired medio-lateral location and screwed back in place. Finally, the optic fiber was placed inside the implant in contact with the coverslip and maintained with a M1 screw.

During data collection, mice could freely behave in the glass corridor. No food or water restriction or reward was used. A typical experimental session consisted of 50 optogenetic stimulations delivered at random intervals (range: 2-18 s). During the experiment, the irradiance at the tip of the fiber and the duration of the stimulation randomly varied between trials in the 7.5-120 mW/mm² and 50-1600 ms ranges respectively. Tens of sessions could be carried in the same animal over several days to ensure a sufficient number of trials for each set of stimulation parameters. Between each session the mouse was kept in its home cage in order to minimize stress.

Movies were collected at 100 frames per second with a spatial resolution of 1200x220 (bottom view camera) and 1200x800 pixels (side view camera). The acquisition software was written in LabVIEW and uses one National Instruments board (NI-PCIe-6353) and a connection block (BNC-2090A, National Instruments) to trigger the optogenetic stimulations while simultaneously recording the movie of the animal.

During EMG recording experiments, EMG signals were acquired at 10 kHz with a 50x custom-made amplifier connected to the National Instruments system. Two small diameter (0.5 mm) wires were used to connect the amplifier to the pins cemented on the implant. The wires were passed through a ring above the arena and ran along the optic fiber in order to minimize mechanical constraints on the animal movement.

### Behavior analysis

All analyses were done using the MATLAB software (Mathworks) and performed offline. Both camera views of the animals were processed by the same algorithm.

### Animal movement quantification

To quantify the movement of the mouse, we computed the frame-to-frame differential movie of the animal. We then applied a threshold on the resulting movie and defined the global movement of the animal as the sum of each frame, i.e., the number of pixels whose values had significantly changed. The threshold was set at mean + 5*SD of the movie noise. The resulting traces were smoothed with a gaussian filter (30 ms width). All traces were normalized by the mean amplitude of M_OFF_ over the whole dataset. The different peaks values and timings were defined as local maxima in a specific temporal window (between 0 and 250 ms for M_ON_ peak, from 0 to 250 ms after stimulation termination for M_OFF_. In Figure 1L, the distributions were split in two based on gaussian fits for low stimulation powers (merged datasets 7.5 and 15 mW/mm²). “Non-failure” of M_ON_ was defined as a M_ON_ peak value exceeding mean + 3*SD of the first gaussian.

### Animal body barycenter extraction

In order to extract the barycenter of the mouse body, each frame of the movie was saturated and binarized using a fixed threshold (100 for an 8-bit grayscale image). Because the tail and paws of the mouse would perturb the measurement, the resulting shape was then eroded using a binary disk (30 pixels radius) to extract the trunk of the animal. The mouse body barycenter was defined as the barycenter of the resulting binary shape. Barycenter trajectories were smoothed using a gaussian filter (30 ms width). In Figure 2E, the posture retrieval was defined as the timing at which Zb (the Z coordinate of the segmented mouse body barycenter) returned to baseline levels (mean ± 3*SD of the Zb values before light stimulation). ΔZbmax was defined as the mean value of ΔZb 30 ms around stimulation termination.

**Figure 2.**
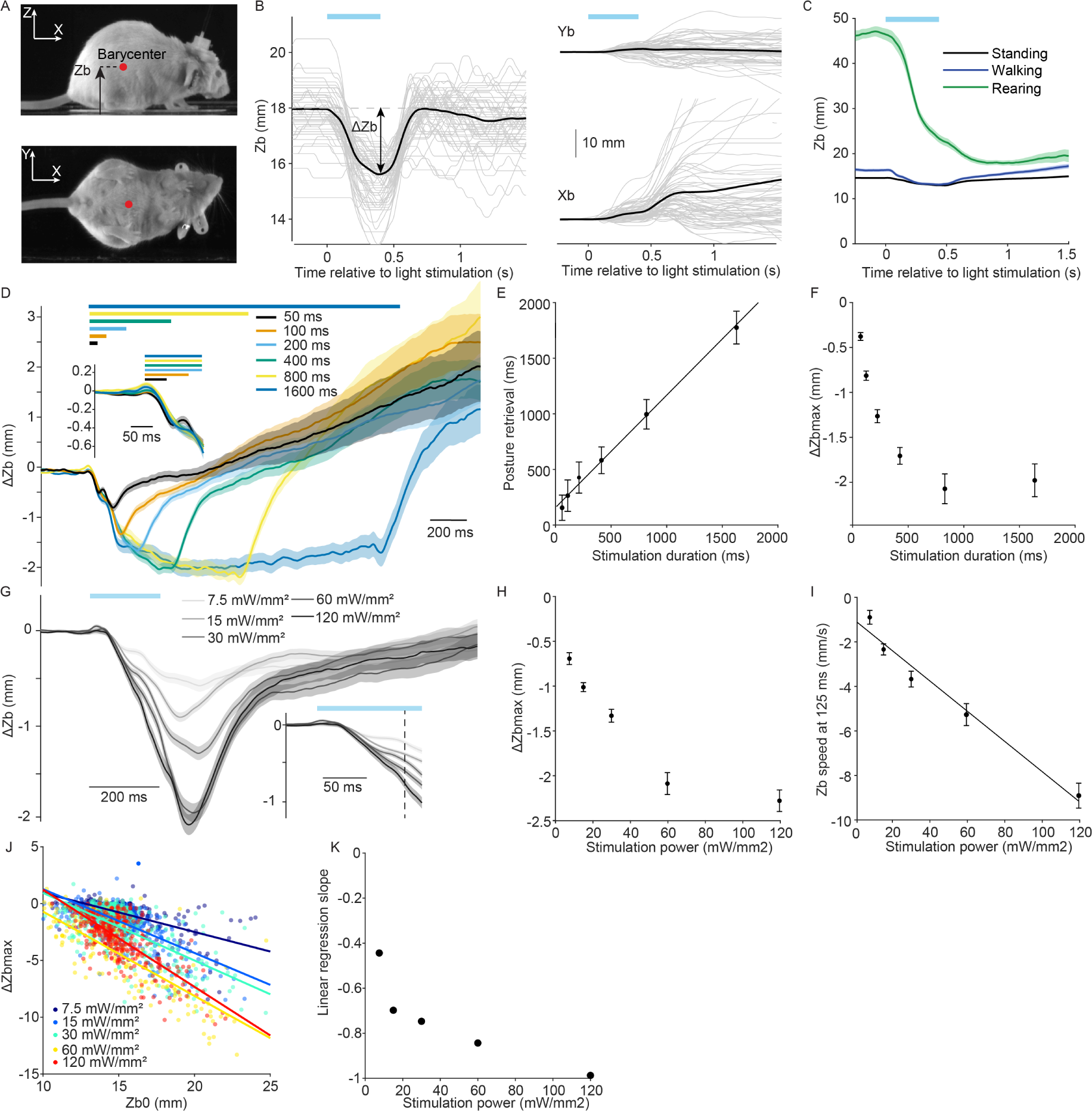
The direct effect of the stimulation is a postural collapse. **A**. The barycenter of the mouse body was computed from two camera views. **B**. Individual and mean barycenter trajectories (gray and black traces respectively) following a 30 mW/mm² 400 ms optogenetic PC stimulation at the lobule IV/V midline (n = 63 trials). **C**. Time course of the Z coordinate of the body barycenter around the optogenetic stimulation for three different behavioral states (see Experimental Procedures). Curves represent mean *±* SEM (n = 8 animals, 30 mW/mm² 400 ms stimulations). **D**. Time course of ΔZb for different stimulation durations (n = 8 mice). Curves represent mean *±* SEM. **E**. Start of posture retrieval plotted against stimulation duration (R = 0.998, y = 1.01x + 168 ms). Data are mean *±* SEM. **F**. ΔZbmax plotted against stimulation duration. **G**. Time course of ΔZb for different stimulation powers (n = 8 mice). Curves represent mean *±* SEM. **H**. ΔZbmax plotted against stimulation power. **I**. Zb speed at stimulation onset (from 25 to 125 ms following stimulation onset) plotted against stimulation power (R = 0.972, y = -6.62e-3x - 0.160). **J**. ΔZbmax plotted against Zb0 (before light stimulation) for individual trials and different stimulation powers (400 ms stimulations). Lines represent the linear regression. **K**. Slope of the linear regressions from **J** plotted against stimulation power.

In Figure 1C and 2C, rearing was defined with a threshold on the mouse barycenter altitude in the 100 ms preceding light stimulation (2.5 cm from the ground). Walking was defined with a threshold on the animal speed in the 100 ms preceding light stimulation (>5 cm/s). The state “standing” was defined by excluding the rearing and walking trials.

### EMG post-processing

The EMG signals were processed with a custom MATLAB routine. EMGs were band-pass filtered with a 4^th^ order Butterworth filter (cut-off frequencies, 50 and 1000 Hz) and the envelope of the signal was computed using a sliding RMS window over 30 ms. The envelope amplitude in Figure 3 was defined as the maximum of the envelope in the M_ON_ window (0-250 ms after light stimulation commenced). Muscle contraction was considered significant when it exceeded mean + 3*SD of the baseline levels (-100 to 0 ms before stimulation). Contraction onset was defined as the earliest timing corresponding to a significant contraction during light stimulation.

**Figure 3.**
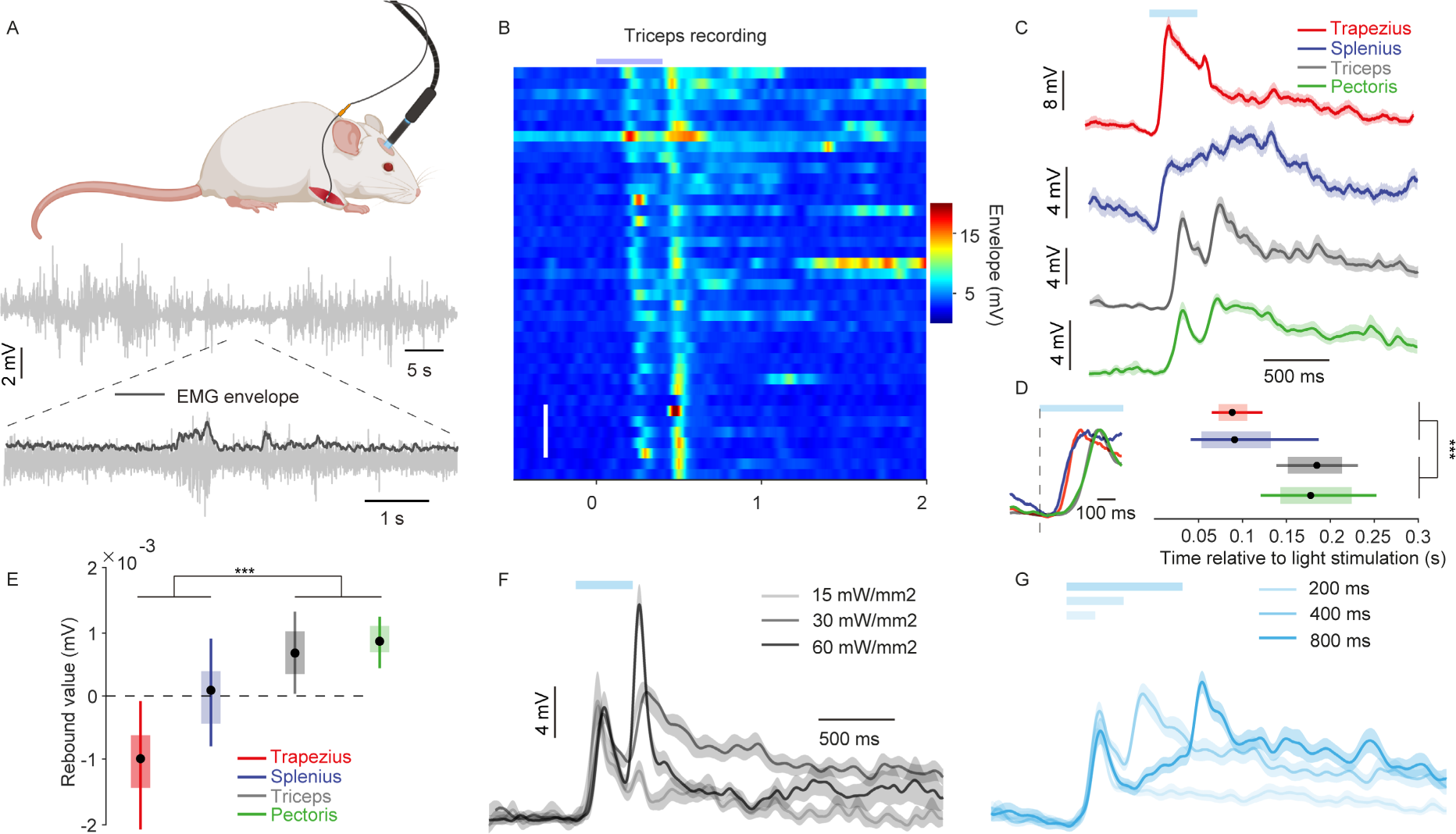
Body muscles involvement during PC optogenetic activation. **A**. (Top) Mice were implanted with EMG electrodes in several body muscles (see Experimental Procedures for details). (Bottom) Raw EMG traces were temporally filtered and processed to compute the envelope of the signal (see Experimental Procedures). **B**. Example raster of the EMG envelope around optogenetic PC activation for the triceps muscle (30 mW/mm² 400 ms stimulations). White bars represent 5 trials. **C.** Average time-course of the EMG envelope for the recorded body muscles (30 mW/mm², 400 ms stimulations, n = 2 mice for the trapezius, 4 mice for the splenius, 3 mice for the triceps). Data are mean *±* SEM. **D**. (Left) Normalized envelopes of the recorded body muscles around stimulation onset. (Right) Muscle contraction onset timing for the body muscles recorded in B. Plain dots represent the median, shaded boxes the 25-75% and colored lines the 5-95% distributions limits respectively. **E**. Relative muscle contractions for the muscles recorded in B following light termination. Same representation as in D (400 ms 30 mW/mm² stimulations). **F**. Evolution of the EMG envelope of the triceps muscle with stimulation power (400 ms stimulations, n = 3 mice). **G**. Evolution of the EMG envelope of the triceps muscle with stimulation duration (30mW/mm² stimulations, n = 3 mice).

### Points of interest tracking and realignment along the animal body axis

Specific landmarks on the animal body were tracked using the DeepLabCut software (Mathis et al. 2018). We tracked both the snout and the base of the tail of the mouse for different purposes.

The tail position was used to realign the movie in the mouse referential. First, a movie centered on the barycenter of the animal was extracted from the original bottom view recording. This movie was processed by the DeepLabCut software for tail tracking (Mathis et al. 2018). Then, the bottom view video was rotated to be aligned with the body axis of the animal, so that on the resulting movie the mouse had a constant orientation. The tail position was also used to horizontally flip the lateral view movie of the mouse in order to keep its body in a constant orientation for all trials. Finally, the resulting movies were again processed through DeepLabCut for snout tracking. The resulting trajectories were smoothed using a gaussian filter (30 ms width). When computing the snout trajectory, we only kept the trials for which its detection was reliable based on the confidence on the detection returned by the algorithm (>0.8).

### Statistics

All statistics were performed using the MATLAB software (Mathworks). Unless otherwise noted, paired-test significativity was assessed with a two-sided Wilcoxon rank sum test and unpaired test significativity with a Wilcoxon signed rank test.

### Trial number

In Figures 1 and 2, each condition comprised between 171 and 513 trials (n = 10 mice). In Figure 3, each condition comprised between 95 and 335 trials (n = 3 mice for the triceps, 2 mice for the trapezius, 4 mice for the splenius and 2 mice for the pectoris). In Figure 4 B, each condition comprises between 132 and 392 trials (n = 8 mice). In Figure 4E-M, each condition comprises between 196 and 258 trials (n = 5 mice).

**Figure 4.**
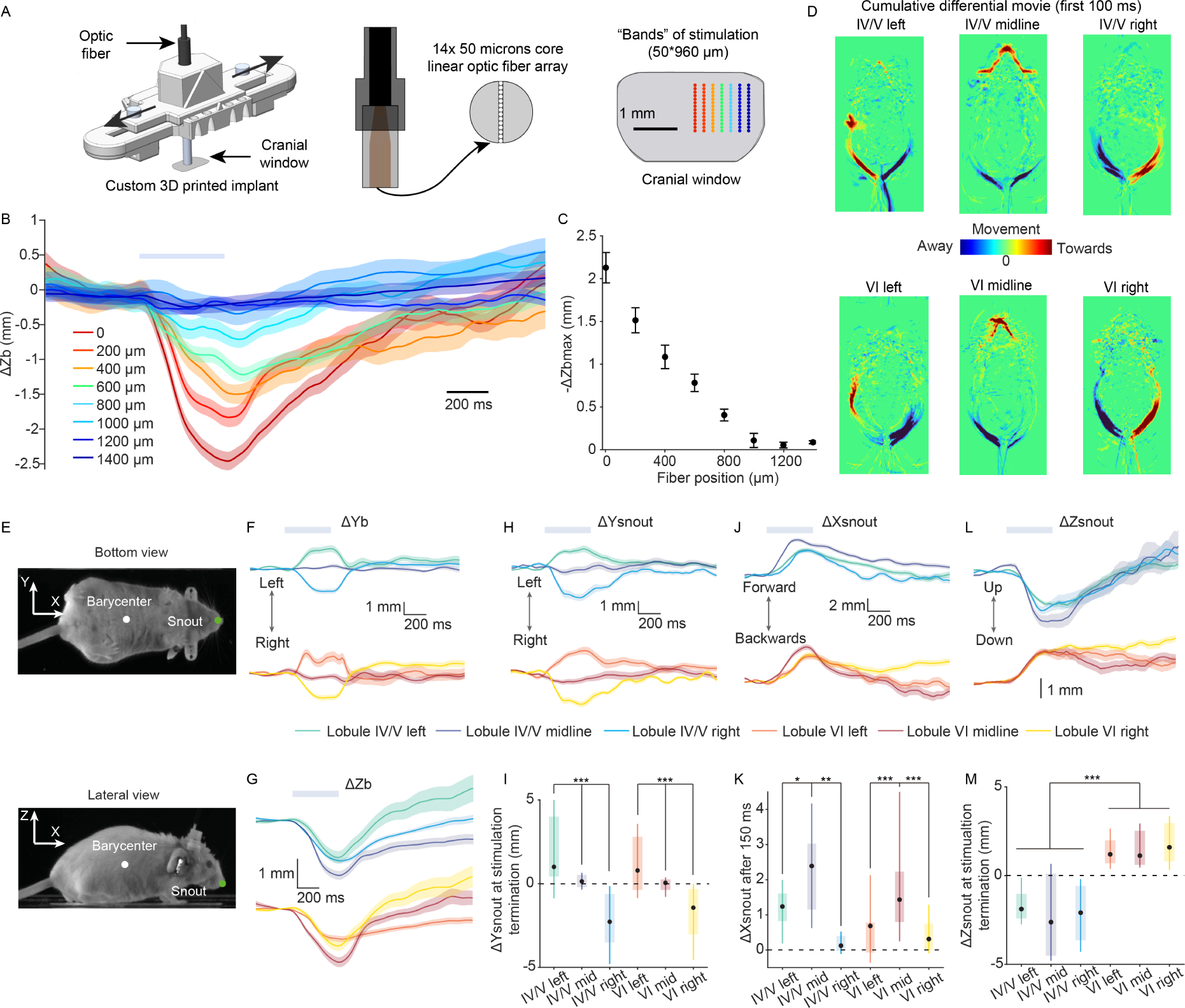
Optogenetic mapping of the anterior vermal output. **A**. (Left) Custom implant used to map the cerebellar vermis. The implant is composed of a top part which can slide on a bottom fixed part. (Middle) The implant is designed to position a custom linear fiber array over the cerebellar cortex. Moving the array allows sequential stimulation of sagittal PC bands with high spatial specificity. **B**. Time course of ΔZb for different medio-lateral locations over the vermal lobule IV/V. The coordinate 0 refers to the midline. The fiber was moved in the rightward direction. Curves are mean *±* SEM (n = 4 mice, 50 trials per mouse, 30 mW/mm² 400 ms stimulations). **C**. ΔZbmax plotted against the medio-lateral location of the optic fiber array. Data are mean *±* SEM. **D**. Average cumulative differential movie between 0 and 100 ms after stimulation commenced for different fiber locations. **E**. After a realignment of the mouse body along the tail-body barycenter axis, the snout of the animal was tracked with both camera views using the DeepLabCut software (see Experimental Procedures). **F**. Time course of the differential ΔYb coordinate of the animal segmented body barycenter for different medio-lateral locations in the lobules IV/V and VI. **G**. Same as F but for ΔZb. **H**. Same as F but for ΔYsnout. **I**. ΔYsnout at stimulation termination for the different fiber locations. Plain black dots represent the median, shaded boxes the 25-75% and colored lines the 5-95% distributions limits respectively. **J**. Same as F but for ΔXsnout. **K.** ΔXsnout 150 ms after light stimulation commenced for the different fiber locations. **L.** Same as F but for ΔZsnout. **M.** Same as H but for ΔZsnout.

## RESULTS

### Optogenetic stimulations of the anterior vermal Purkinje cells elicit a movement sequence

To investigate the role of the cerebellar vermis in postural control, we used a transgenic mouse model expressing channelrhodopsin-2(H134R) uniformly and specifically in PCs (Chaumont et al. 2013). We implanted these mice with a small-core optic fiber (200 µm), which allowed for confined light stimulations of PCs in the freely moving animal. The mice were introduced in a transparent glass arena where they could behave naturally (Figure 1A), thus preventing potential confounding factors encountered in constrained animals. The animal was monitored from the side and the bottom at high frame rate (100 Hz) to allow for a precise 3D spatio-temporal movement analysis.

When optogenetic stimulations were delivered at random intervals to the midline of the lobule IV/V, we observed that a 400 ms stimulation at 30 mW/mm² always elicited a movement (Movies S1 and S2). By computing the frame-to-frame difference of the movie (100Hz), which is a good proxy for the total quantity of movement of the animal, we were able to dissect this movement’s dynamics (Figure 1B). The result revealed two peaks of movement (Figure 1C), which were present across behavioral states. The first (M_ON_) lagged the light stimulation by 163 ± 34 ms (mean ± SD) and the second (M_OFF_) occurred once the light stimulation was terminated. When the animal was rearing on its hind legs, M_ON_ was in registration with the animal’s collapse, while at rest on all four legs, or walking, M_ON_ corresponded to a forward acceleration and a forward step (Movie S2). To exclude confounding factors linked to an ongoing movement we limited our analysis to movements from stimulations of animals that were “at rest” and standing on their four legs (see Experimental Procedures).

Varying the duration of the optogenetic stimulation confirmed that M_OFF_ was indeed time-locked to the OFF of light stimulation (Figure 1D-E, R = 0.999, p = 4.5e-9). For all durations of stimulation, we observed the same stimulation latency of M_ON_ (Figure 1D and 1F), except for stimulations under 200 ms, for which M_ON_ and M_OFF_ appeared to overlap. For 50 ms duration stimulations M_OFF_ would fall within the M_ON_ latency window. Indeed, a single movement occurred and with a shorter latency than normal M_ON_ (Figure 1D, 145 ± 29.8 for 50 ms stimulations vs M_ON_ latency of 164 ± 32.3 ms for stimulations longer than 200 ms, p = 2.7e-17, Wilcoxon rank sum test). Thus, the M_OFF_ is not a correction of M_ON_ (the postural defect caused by the stimulation), but is directly caused by stimulation cessation. The amplitude of M_OFF_ increased with stimulation power (Figure 1G-H), but decreased with stimulation duration (Figure 1I). This is consistent with the idea of a movement provoked by rebound firing in the DCN when PC inhibition resumes to its normal level, which may decrease with stimulation time due to DCN cells adaptation (Witter et al. 2013; Lee et al. 2015). To further dissect the effect of the optogenetic Purkinje cells stimulation, we thereon focused on long stimulations (200-1600 ms), for which M_ON_ was fully developed, did not overlap with the M_OFF_ and had a constant amplitude (Figure 1D and 1J).

Although M_ON_ seemed to increase and saturate with stimulation power (Figure 1D and 1K), closer examination of the response at sub-saturating stimulations power (7,5 and 15 mW/mm²) revealed that it did not occur for every stimulation. In fact, failures resulting in a first mode of amplitude distribution were observed, while full-blown responses had similar amplitudes to those observed at high stimulation power (Figure 1L). The failure probability of M_ON_ gradually decreased towards zero for high stimulation powers (Figure 1M). When we corrected for this factor, the non-failure amplitude was independent from stimulation power (Figure 1K, p = 0.67 between 15 mW/mm² and 120 mW/mm², Wilcoxon rank sum test), thus qualifying for a reflex movement. Movie inspection revealed that M_ON_ generally corresponded to a forward step (Movie S2).

Previous optogenetic stimulation of vermal PCs performed in head-fixed animals was shown to evoke a movement exclusively when light stimulation terminated (Witter et al. 2013). While we also observe this light termination effect in a freely moving animal, we additionally report the occurrence of a M_ON_. The reflexive nature of this movement raises the possibility that it does not represent the direct effect of the optogenetic stimulation but a secondary consequence of the postural defect through the optogenetic PC stimulation. This led us to analyze in more detail the displacement of the animal body in the period immediately following light stimulation and preceding M_ON_.

### The direct effect of the stimulation is a postural collapse

In order to better characterize the animal movement kinematics, we segmented the mouse, recording simultaneously the body from side and bottom cameras, and extracted the coordinates of the body barycenter, which approximates its center of gravity (Figure 2A, see Experimental Procedures). Surprisingly we discovered that during the light stimulation the segmented mouse body undergoes a striking decrease of the altitude of the barycenter (Zb) which immediately follows light onset (Figure 2B). This effect on the mouse barycenter was observed in all initial postural conditions (Figure 2C). The altitude drop robustly preceded M_ON_ in 85 % of the trials (429/504 trials, 74 ± 38 ms for Zb decrease versus 124 ± 47 for Xb, p = 1.4e-59, Wilcoxon signed rank test). The downward movement of the body therefore likely constitutes the primary effect of our optogenetic PC stimulation, hereafter referred to as M_ON-DIRECT,_ while the M_ON_ forward step would constitute a compensatory reflex, hereafter referred to as M_ON-REFLEX_. To test this interpretation, we varied the stimulation parameters and found that with increasing stimulation duration (at fixed power) the body altitude-collapse gradually developed, and then saturated to a constant altitude during long stimulations. Collapse always started to recover immediately after stimulation termination (Figures 2D and 2E), again suggesting that M_ON-DIRECT_ is the primary and direct consequence of increased PC activity. Varying the power of a fixed short-duration light stimulation (200 ms) led to a graded postural collapse which saturated for high intensities (Figures 2G and 2H). Strikingly, the vertical speed of the barycenter at the onset of the collapse (from 25 to 125 ms after commencing stimulation) was linearly correlated to the stimulation intensity (Figure 2G (insert) and 2I), indicating an imbalance in anti-gravity muscle tone proportional to the optogenetic stimulation.

We then looked, in greater detail, at the trial to trial variability in the movement induced from light stimulation. At all stimulation powers the loss of barycenter altitude at steady-state was correlated with the initial altitude of the mouse body (Figure 2J). As the power increased, however, the slope of the linear regression increased and converged towards unity (Figure 2J and K, from -0.44 for 7.5 mW/mm² stimulations to -0.99 for 120 mW/mm² stimulations). At these saturating powers the final loss of barycenter altitude is therefore constant and corresponds to the lowest position that can be reached, essentially when the ventral surface of the mouse body rests on the surface of the arena. At non-saturating powers the stimulation seemed to impose a multiplicative gain on the maintained antigravity muscle-tone (which sets the altitude of the animal), leading to larger falls for those postures where the animal’s body was being held farther off the surface of the arena. Part of this effect could arise from the fact that the final position of the barycenter is determined by the fraction of the animal’s weight which is already in contact with the ground and supported by it, thus counterbalancing the optogenetic muscle tone loss.

Taken together, these results show that the direct effect of the light stimulation was a loss of anti-gravity maintenance. Overall, the stimulation effect can be decomposed into four phases (Movie S3). First, an anti-gravity postural collapse starts about 40 ms (38 ± 15 ms for a 30 mW/mm² 400 ms stimulation) after the light stimulation commences, M_ON-DIRECT_. Second, at 120 ms post stimulation, a compensatory reflex engages, M_ON-REFLEX_, generally a step forward, aiming at restoring balance. Third, the animal undergoes a loss of anti-gravity muscle tone, which is prolonged even while the animal attempts to execute voluntary forward steps during the stimulation. Finally, after termination of the optogenetic stimulation, the mouse performs a phasic motor response, M_OFF_, most likely triggered by a rebound in the deep cerebellar nuclei, while slowly retrieving its postural altitude.

### Body parts are differentially involved during optogenetic PC activation

We sought to investigate the involvement of different body muscles in the postural perturbation evoked by the optogenetic stimulations. To address this, we implanted EMG electrodes in two neck muscles involved in upward head movement (trapezius and splenius), and two muscles involved in body support by the forelimbs (triceps and pectoris), and extracted their muscle tone from the recorded signal by computing the EMG envelope (Figure 3A, see Experimental Procedures). During the optogenetic stimulation, we observed that the movement of the mouse was consistently associated with contractions of all four of the recorded muscles (Figure 3B), and all started to contract at least 100 ms after the stimulation commenced. Therefore, we considered these muscle contractions a signature of the mouse postural reflex. As expected, contractions of these muscles were also observed at posture retrieval (Figure 3B and 3C).

Interestingly, based on the timings of contraction after stimulation commenced, we observed that the trapezius and splenius muscles contracted significantly before the triceps and pectoris. As shown in Figure 3D, the recordings revealed trapezius and splenius starting to contract at 99 ± 38 ms and 112 ± 59 ms respectively (p = 0.58, Wilcoxon rank sum test). Then, at 179 ± 57 ms the pectoris begins to contract, and finally the triceps (183 ± 45 ms for the triceps, p = 6.4e-36 between the two muscle pairs of trapezius & splenius versus pectoris & triceps). On a trial-to-trial basis the trapezius and triceps muscle contractions were highly correlated to M_ON-REFLEX_ amplitude (R = 0.93 and 0,65, p = 5.3e-30 and 1.6e-19 respectively), confirming their involvement in this phase of the movement. Correlations to the two other muscles were weaker albeit significant (R = 0.37 and 0.27, p = 4.5e-8 and 4.72e-4 for the pectoris and splenius respectively). Lower significance could be explained by the smaller impact of these muscles on the global body position, as quantified by the cameras. Thus, the muscles could be placed in two groups. Moreover, we observed that, similarly to the M_ON-REFLEX_ amplitude, the amplitude of the M_ON-REFLEX_ contraction was independent from both the stimulation power (Figure 3E, triceps muscle, 400 ms stimulations, n = 3 mice) and duration (Figure 3F, triceps muscle, 30 mW/mm² stimulations). This provides further confirmation that the optogenetic stimulation of Purkinje cells evokes a compound sequential reflex aimed at stabilizing the head and the trunk of the animal. Moreover, in all the recorded muscles, in particular the splenius, the contraction appeared to be sustained during the stimulation (Figure 3B). This indicates that after the postural reflex the mouse tried to compensate for the still ongoing loss of muscle-tone but despite prolonged contractions could not overcome the effect of the perturbation.

Although the EMG did not reveal any significant muscle relaxations in the 120 ms period during body collapse (the M_ON-DIRECT_ period), we reasoned that muscles which were perturbed when stimulation commenced ought to display a clear rebound of activity upon termination of stimulation. We indeed observed such a rebound in pectoris and triceps, which are both muscles that act to maintain posture by counteracting gravity (Figure 3G, rebound values 0.54 ± 0.51, 0.59 ± 0.22 for triceps and pectoris respectively). In contrast, we did not see a similar rebound in the two neck muscles (Figure 3G, rebound values -0.81 ± 0.20 mV, 0.04 ± 0.28 for the trapezius and splenius respectively). For the triceps, recordings were stable enough to verify that the light-OFF response was graded by the stimulation power and duration (Figures 3E-F). Accordingly, we ascribe these pectoris and triceps contractions to the M_OFF_ rebound activity originating from the DCN. In conclusion, pectoris and triceps are two muscles likely directly affected by the optogenetic PC stimulation.

### Optogenetic mapping of the anterior cerebellar vermal output

We next set out to map the output of the anterior vermis. For this, we produced a 3D-printed implant for mounting a custom optic fiber array (14 cores of 50 µm aligned on a linear array) over a cranial window that spanned the whole medio-lateral extent of the anterior vermal cerebellum. Thus, sequential optical stimulations of mild intensity (10 mW/mm², 400 ms stimulation) could be delivered to narrow sagittal bands of PC (Figure 4A) at different medio-lateral locations in the cerebellar lobule IV/V. As we gradually moved the fiber from a medial to lateral locations, we observed that the postural collapse amplitude progressively decayed and disappeared once outside of the vermis (Figure 4B-C, n = 4 mice). The mid-effect was found around 500 µm laterally, which is in accordance with a broad (wider than a cortical microzones) module of DCN cells previously defined by genetic profiling, PC axonal projections to the DCN and zebrin immunolabeling patterns (Hawkes, Colonnier, and Leclerc 1985; Sugihara et al. 2009; Voogd 2011; Fujita, Kodama, and Du Lac 2020).

In order to assess the existence of functional subregions, we refined our analysis of the evoked movement and used machine learning to track points of interest on the animal’s body. The base of the mouse tail was tracked, allowing us to realign its body around the body axis (see Experimental Procedures). We then computed the average cumulative differential movie of the animal between 0 to 100 ms after stimulation commenced (before M_ON-REFLEX_, the postural reflex), which produced a body map of the mouse movement over this period (Figure 4D, 400 ms stimulations, n = 3 mice). Comparing these body maps for different stimulation locations, we found that the mouse movement was lateralized in both lobule IV/V and VI (Figure 4D), which was also observed when measuring the Y coordinates of the mouse body barycenter (Figure 4E and F, ΔYb body lateral displacement of 2.1 ± 1.61 mm and -2.33 ± 1.26 mm for left and right lobule IV/V stimulations respectively, 1.22 ± 1.63 mm and -1.8 ± 1.14 mm in lobule VI, p = 1.4e-20 and 8.4e-15 in lobule IV/V and VI respectively), the mouse body falling ipsilateral to the side of stimulation. In contrast, a straight downward movement of the animal body was seen for midline stimulations (Figure 4D-F, 0.19 ± 0.58 mm and -0.25 ± 0.74 mm ΔZb body vertical displacement in lobule IV/V and VI respectively). As expected, a global body collapse was observed at all locations (Figure 4G, bottom panels).

The mouse’s snout is a good proxy of head movement (Figure 4E, see Experimental Procedures). While the Y coordinate of the snout reported a lateralization of the stimulation effect (Figure 4H and I, average ΔYsnout at stimulation termination of 1.91 ± 1.32 mm and - 3.1 ± 1.69 mm for left and right lobule IV/V stimulations respectively, 1.46 ± 0.99 mm and -2.56 ± 1.25 mm in lobule VI, p = 4.0e-16 and 2.3e-19 in lobule IV/V and VI respectively), the animal’s snout also displayed an initial additional forward movement for medial stimulations compared to lateral stimulations (Figure 4J and K, ΔXsnout 150 ms after light onset 2.52 ± 1.25 mm and 1.52 ± 1.34 mm for medial vs lateral stimulations in lobule IV/V, 2.47 ± 1.63 mm and 1.50 ± 1.81 mm in lobule VI, p = 3.3e-4 and 9.7e-3 in lobules IV/V and VI respectively). Stimulations in lobule IV/V produced similar snout (Figure 4L) and body barycenter altitude decrease (Figure 4G). In contrast though, stimulations in lobule VI evoked an increase in the snout altitude during the stimulation (Figure 4L and M, average ΔZsnout at stimulation termination -2.04 ± 1.11 mm in lobule IV/V vs 1.53 ± 0.80 mm in lobule VI, p = 1.4e-85 between the two lobules), although the mouse experienced a postural collapse of the body barycenter (Figure 4G), suggesting a contraction of the dorsal neck muscles. These results indicate that a fine-grain functional somatotopy of the anterior vermis within a genetically and immunohistochemically identified module.

## DISCUSSION

Our results from optogenetic stimulations of cerebellar PCs argue for a direct involvement of the anterior vermis in anti-gravity postural maintenance: we find that a postural collapse is the first event to occur upon sustained increased firing of PC and is directly modulated by stimulation parameters (Figure 2). While lesions and tracing studies have long pointed to a role of the vermis in basic postural functions (Chambers and Sprague 1955a; 1955b; Joyal et al. 1996; Manni and Petrosini 2004), this is to our knowledge the first direct experimental evidence in the intact freely behaving animal of a vermal role in postural maintenance.

### Anterior vermis involvement in anti-gravity postural maintenance

Previous studies had intended to assess the effects of optogenetic manipulation of PC activity in the dorsal vermis of the mouse (Witter et al. 2013; Hoogland et al. 2015; Heffley et al. 2018). One of them described a behavioral response of the animal exclusively at the termination of PC stimulation, which was characterized by whole-body twitches (Witter et al. 2013). We similarly observed a movement at stimulation termination, which not only involves a posture retrieval of the mouse but also a rebound movement graded by stimulation power (Figure 1H), which we further characterize as a rebound contraction in specific muscles (Figure 3E-G). Witter et al. performed experiments in head-fixed animals, however, which most likely prevented the observation of the anti-gravity effect reported here. Interestingly, a subsequent report using head-fixed animals placed on a rotating disk (Hoogland et al. 2015) pointed to an initiation of stepping around 200 ms after stimulation commenced. This timing arguably corresponds to the compensatory postural adjustment we observe in this study, which often manifests as a forward step. The same study reported a slow-down of stepping when the animal had already been walking, while in contrast we observed a continuation of the ongoing forward movement (Figure 2B). These observations can be reconciled if one accepts the plausible argument that in head-fixed animals (Hoogland et al. 2015), the stimulation will cause a weakening of the forelimbs and thus reduce stepping without causing an imbalance of the body towards the front. In freely-moving animals (as in this report), reduced anti-gravity tone of the forelimb causes a downward movement of the head and a loss of equilibrium, leading to a strong maintenance reflex, hence a stepping forward. Thus, previous reports are consistent with our results but appear to have failed to uncover the direct effect of the stimulation, presumably because of the experimental constraints encountered in a head-fixed animal.

### Compensatory postural reflex elicited by optogenetic stimulation

We identified a compensatory postural reflex following postural collapse of the animal body, which is associated with a sequence of muscle contractions. The first pair of muscles to contract (trapezius and splenius) are neck muscles whose role is to maintain the head of the animal up. The contraction of these muscles 100 ms after optogenetic stimulation commences matches a short slowing in the animal’s barycenter drop in altitude (Figure 2D), indicating a first reflexive component. In contrast, the second set of muscles (pectoris and triceps) that contract 70 ms later are more likely involved in the subsequent compensatory step performed by the animal, which translates in strong correlations with this part of the movement. Moreover, the reliable contraction observed in these two muscles at stimulation termination (Figures 3G) indicates that they are likely the ones directly affected by the stimulation.

The compensatory reflex may involve several extracerebellar brain regions at the spinal and brainstem levels, in particular in link with the vestibular system (Horak 2006). However, in particular for the muscles directly affected by the stimulation, the reflex could have a cerebellar origin. It is known that PCs can control their own afferent climbing fiber discharge with a minimal delay of 80-100 ms (Chaumont et al. 2013). Therefore, we can expect a synchronous CF discharge in the stimulated PCs around the timing of reflex initiation, which would participate in the reported reflexive movement.

Interestingly, we observed that muscle contractions were often prolonged during the whole stimulation duration (Figure 3B-C). This signals a prolonged compensation of the postural loss, which translates in a prolonged forward movement of the animal, in particular at long durations (Figure 1D). This compensation is nonetheless insufficient to outweigh the stimulation effect, since the barycenter altitude only recovers after stimulation termination (Figure 2D-E). Thus, the cerebellum does not decrease a specific drive associated to anti-gravity function but exerts a global gain function on muscle tone, which prevents these muscles from being properly used in the subsequent motor command generating the forward stepping during the optogenetic stimulation.

### Influence of PC activation on muscle tone

Because the anterior vermis has been associated with the anterior axial part of the body (Chambers and Sprague 1955a; 1955b), we targeted axial and proximal muscles in the neck region for our EMG recordings during optogenetic stimulations. Surprisingly, we did not observe any muscle relaxation in our EMG recordings (Figure 3B-C). While the affected muscles could belong to other body regions, such as the back or the belly of the animal, the specificity of the contraction at stimulus termination in two muscles strongly indicates that they could be amongst the ones controlled by the vermal lobule IV/V at midline. This hypothesis would fall in line with the known representation of the forelimbs in the vermis (Heffley et al. 2018; Wagner et al. 2021). The absence of muscle tone loss in our recordings could be explained by experimental limitations. In Figure 2I, we observe that the mouse barycenter decreased 1 mm 100 ms after stimulation commenced. We can thus calculate that the acceleration of the animal’s body is around 0.1 m/s^-2^, which roughly represents a hundredth of earth gravity (9.8m/s^-2^). Therefore only a few percent of muscle tone loss could be sufficient to trigger the observed effect. If these calculations are correct, the resulting EMG variation would be difficult to pick up in our recordings.

### Cerebellar output activity during PC stimulation

Although we did not record the PC activity during our experiments, based on previous studies, we can confidently infer that our PC optogenetic stimulations elicit a graded DCN inhibition (Chaumont et al. 2013; Witter et al. 2013). This assessment is non-trivial because strong PC stimulations have been shown to induce a depolarizing block in PC activity, which could result in net DCN disinhibition (Chaumont et al. 2013). The stimulation power we report was measured directly at the fiber tip. Because a glass coverslip is positioned between the fiber and brain, we can extrapolate based on our fiber NA (0.22) and coverslip thickness (150 μm) that there is a 1.5-fold increase in the illuminated surface when the light reaches the brain. The irradiance is therefore divided by 1.5, which yields a range of 5-90 mW/mm² at the brain surface, which is similar to that used in the study characterizing the mouse line we used (Chaumont et al. 2013). Moreover, even if at high power stimulations PCs directly below the fiber tip were to undergo a depolarizing block, the diffusion of light in living tissue with increasing power would recruit large numbers of PCs around the stimulation site and globally result in a net DCN inhibition (Chaumont et al. 2013). Finally, the postural collapse we observed was consistent over the entire ranges of durations and powers assayed in our experiments. It is therefore most probable that, even at high intensities, we effectively inhibit the cerebellar output from the DCN instead of activating it.

### Spatial specificity of the stimulation

Our optogenetic mapping of the vermis relies on spatial specificity of PC excitation. In order to minimize the extent of the stimulated area, we used a custom optic fiber linear array composed of small-core 50 µm optic fibers, positioned over the vermis in a parasagittal orientation (Figure 4A). It must be noted, however, that beam divergence through the cranial window as well as diffusion inside brain tissue likely increase the extent of the stimulated area. Given the numerical aperture of the optic fiber (NA = 0.22), the glass index (1.5) and the glass window thickness (150 µm), we can expect a divergence through the thickness of the cranial window of:

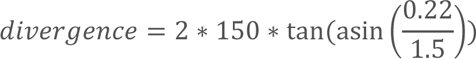

This gives a beam expansion of ∼45 µm in diameter through the glass window, which results in a ∼100 µm width of illuminated area at the cortical surface. Previous studies investigating light diffusion in cerebellar tissues have shown that the diffusion process is predominantly axial (Chaumont et al. 2013; Gysbrechts et al. 2016; Yona et al. 2016), with a 1% isoline located laterally at ∼75 µm from the origin for a 50 µm core optic fiber and at 400 µm in depth. Therefore, we can make the conservative assumption that our stimulation covers a surface of ∼1000*150 µm of cerebellar cortical tissue, which is in the range of the known extent of cerebellar microzones (Kostadinov, Beau, Pozo, et al. 2019) and would correspond to roughly 250 PCs. Thus, our custom optic fiber array allows us to stimulate the vermal PC output at the microzonal scale. Based on previous experiments, we can affirm that PCs more than 200 µm from the stimulation site are very unlikely to be affected (Chaumont et al. 2013). Moreover, the observed lateralization of the postural collapse argues for a degree of spatial specificity in our stimulations, and the ipsilateral effect on the animal body is consistent with the known cerebellar projections to spinal cord motoneurons (Teune et al. 2000).

### Functional organization of the vermal output

The olivary input to PCs is organized in translobular parasagittal bands of CF projections with specific receptive fields (Andersson and Oscarsson 1978), but parallel fibers (PF) distribute sensori-motor information to PCs in the medio-lateral direction (Heck, Thach, and Keating 2007). A long-standing hypothesis, as identified for eyelid closure, ocular saccades and for limb movement (Hesslow 1994; Mostofi et al. 2010; Heiney et al. 2014; Herzfeld et al. 2015; Heffley et al. 2018; Sedaghat-Nejad et al. 2022), posits that the functional unit of the cerebellar cortex, controlling a specific motor function, consists of a small intersectional domain of PCs sharing common CF and PF information. In addition, microzones of correlated CF activity have been identified in relation to various other tasks but direct control of a specific motor output by the corresponding PCs has not been demonstrated in these cases (Mukamel, Nimmerjahn, and Schnitzer 2009; Kostadinov, Beau, Pozo, et al. 2019; Tsutsumi et al. 2019; 2020). This intersectional domain hypothesis is, however, consistent with the known PC-DCN convergence pattern (R. Apps and Garwicz 2000; Voogd 2011) and the known organization of the DCN (Heiney, Wojaczynski, and Medina 2021). In this study, we used an optic fiber with a custom longitudinal small-core array to map the cerebellar output within a specific lobule with fine medial-lateral resolution. The postural function we identified was found to be encoded at the scale of a large vermal zone, roughly corresponding to the A zone of the cerebellar cortex (Richard Apps et al. 2018) and a posturo-motor module identified genetically (Fujita, Kodama, and Du Lac 2020). We show through refined movement analysis that neighboring stimulation sites within this module are associated with different motor outcomes that are putatively related to muscle groups directly controlled by PCs at each stimulation location. This patterning is both medio-lateral and antero-posterior, in agreement with the intersectional domain hypothesis. Interestingly, our posterior stimulation of lobule VI, which may overlap with an output module involved in head and eye orienting movement (Fujita, Kodama, and Du Lac 2020), evoked an upward movement of the head.

Our results therefore establish for the first time a functional microzonal organization of the vermal PC output, and pave the way for future studies aiming at dissecting further the cerebellar functional output in other paradigms.

## AUTHOR CONTRIBUTION STATEMENT

A.G., V.V., J.B. and S.D. designed the study; A.G. performed the experiments; A.G. analyzed the data; A.G. and S.D. wrote the manuscript; J.B. and V.V. critically revised the manuscript. B.M. helped with the design and the assembly of the experimental setup, including user interface programming. A.A. managed mice colonies. All authors have read and approved the final manuscript.

## FUNDING

This project was supported by AMX scholarship from École polytechnique (A.G.), Labex Memolife scholarship (A.G.), the ANR (GluBrain3A ANR 17-CE-16-0014 and FucPoc ANR 20-CE16-0026) and a QLife supporting program (DION-2021).

## DATA AND CODE STATEMENT

Data shown in figures, example raw recordings and MATLAB codes used to analyze the raw data and generate the figures are available at: https://figshare.com/projects/MATLAB_codes_and_data_for_Optogenetic_mapping_of_the_postural_function_in_the_cerebellar_anterior_vermis/180511

## DECLARATIONS OF INTEREST

S.D. and B.M. have ownership shares in Karthala Systems, a commercial supplier of RAMP microscopes.

## ACKNOWLEDGEMENTS

We would like to thank Yvon Cabirou and Gerard Paresys for custom mechanical production and EMG amplifier. We also acknowledge the assistance of the IBENS Fablab (Fédération pour la Recherche sur le Cerveau – Rotary International France 2018) and Victor Llobet in 3D printing procedures. We thank Caroline Mailes-Hamon, the IBENS PFL2 platform and IBENS animal facility. We also thank Astou Tangara and the IBENS Imaging platform for assistance in experimental setup assembly. We thank Laurent Gallais for production of custom laser-cut coverslips.

## Notes

https://figshare.com/projects/MATLAB_codes_and_data_for_Optogenetic_mapping_of_the_postural_function_in_the_cerebellar_anterior_vermis/180511

### Competing Interest Statement

Stephane Dieudonne and Benjamin Mathieu have ownership shares in Karthala Systems, a commercial supplier of RAMP microscopes.

