## Supplementary movies S1-3 for "Identification and organization of a postural anti-gravity module in the cerebellar vermis"

### Slide 1
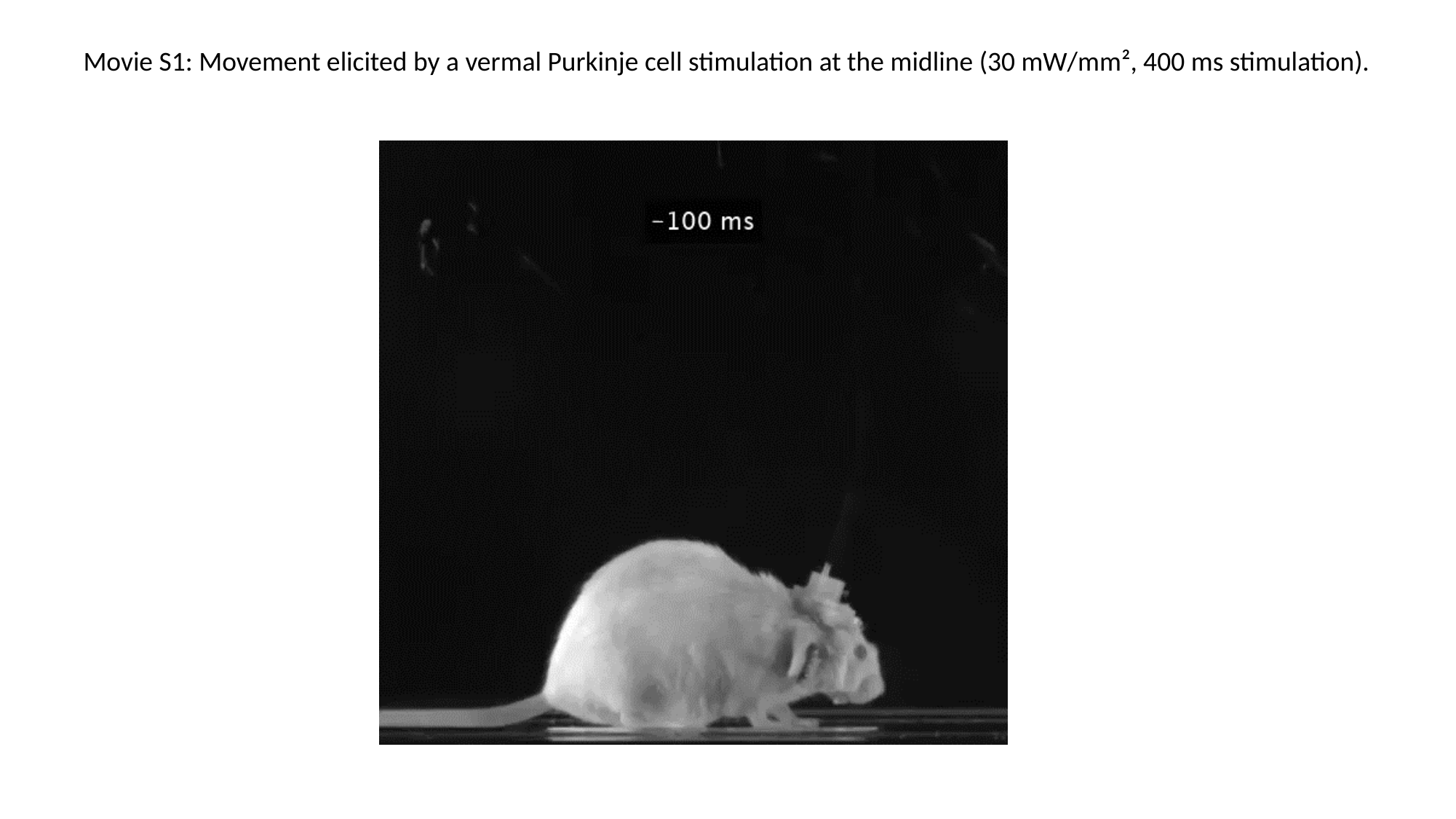

Movie S1: Movement elicited by a vermal Purkinje cell stimulation at the midline (30 mW/mm², 400 ms stimulation).

### Slide 2
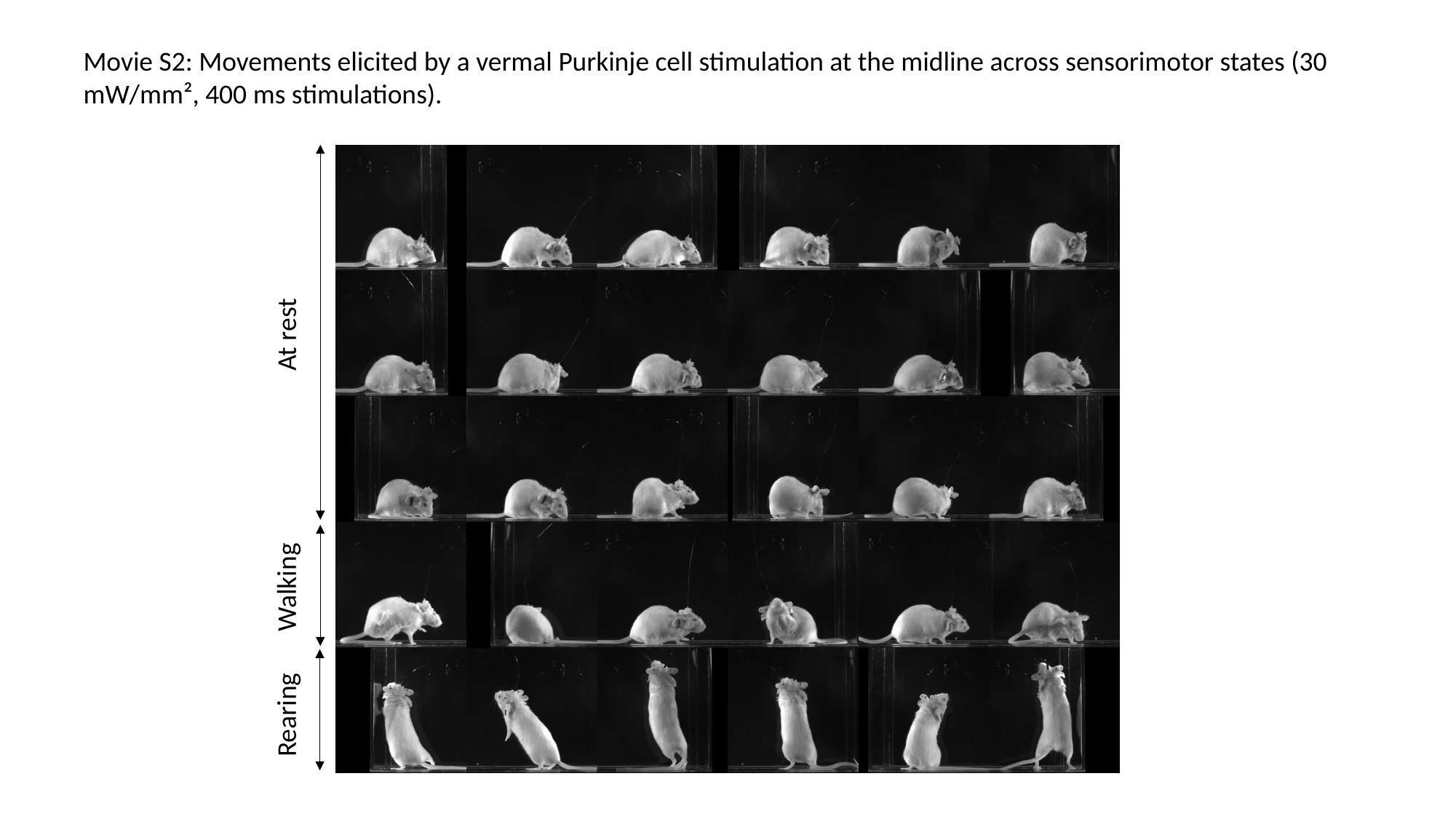

Movie S2: Movements elicited by a vermal Purkinje cell stimulation at the midline across sensorimotor states (30 mW/mm², 400 ms stimulations).
At rest
Walking
Rearing

### Slide 3
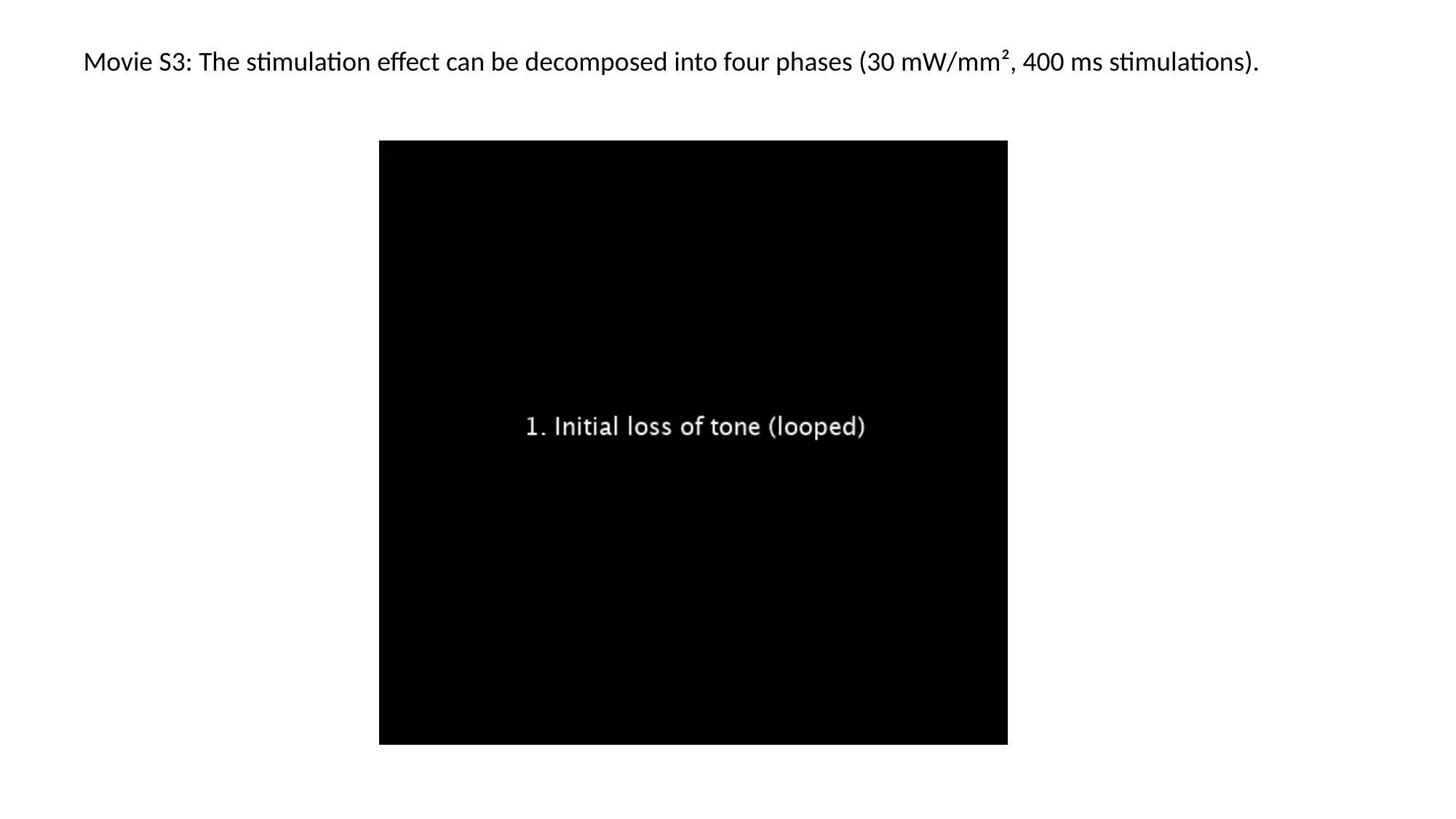

Movie S3: The stimulation effect can be decomposed into four phases (30 mW/mm², 400 ms stimulations).
